# Resilience of BST-2/Tetherin structure to single amino acid substitutions

**DOI:** 10.1101/479402

**Authors:** Ian R. Roy, Camden K. Sutton, Christopher E. Berndsen

## Abstract

Human Tetherin, also known as BST-2 or CD317, is a dimeric, extracellular membrane-bound protein that consists of N and C terminal membrane anchors connected by an extracellular domain. BST-2 is involved in binding enveloped viruses, such as HIV, and inhibiting viral release in addition to a role in NF-kB signaling. Viral tethering by Tetherin can be disrupted by the interaction with Vpu in HIV-1 in addition to other viral proteins. The structural mechanism of Tetherin function is not clear and the effects of human Tetherin mutations identified by sequencing consortiums are not known. To address this gap in the knowledge, we used data from the Ensembl database to construct and model known human missense mutations within the ectodomain to investigate how the structure of the ectodomain influences function. From the data, we identified an island of sequence stability within the ectodomain, which corresponds to functionally or structurally important region identified in previous biochemical and biophysical studies. Additionally, most mutations have little effect on structure, suggesting that they would not affect function. These findings are in agreement with biochemical and cellular studies which suggest that mutations that do not disrupt the alpha helices of Tetherin have little apparent effect on function. Thus, Tetherin sequence is likely less important than structure and this apparent flexibility may allow for greater anti-viral activities with a larger number of viruses.

## INTRODUCTION

Tetherin (BST-2, CD-317) is a membrane bound protein involved in a non-specific, immune response to enveloped viruses, such as HIV-1 and Ebola [Neil, 2013, Sauter, 2014]. During viral budding, one of the transmembrane anchors, usually the C-terminal anchor, becomes embedded in the virion particle membrane and prevents the diffusion of the virion away from the host cell [Perez-Caballero et al., 2009, Venkatesh and Bieniasz, 2013]. Viruses have developed numerous methods to evade viral Tetherin and this is an area of intense study [Sauter, 2014, McNatt et al., 2013]. Additionally, Tetherin is hypothesized to cause an inflammatory response via activation of the NF-kB pathway and may be involved in cell migration [Hotter et al., 2013]. The structural basis for these functions is not fully clear.

Structurally, Tetherin is a homo-dimer consisting of a N-terminal transmembrane helix and a C-terminal membrane anchor bridged by an alpha helical ectodomain (Figure 1) [Andrew et al., 2011]. There are three disulfide bonds within the N-terminal portion of the ectodomain that connect to the two monomers of the dimer and appear to enhance the tensile strength of the protein [Du Pont et al., 2016]. Computational simulation of viral tethering has suggested sequence encoded weaknesses in the C-terminal part of ectodomain that enable bending and flexing of the ectodomain during the transition from membrane bound to bridging the virus and host cell [Ozcan and Berndsen, 2017]. Despite these specific features, upwards of half of the coiled-coil region can be removed without significant loss of function [Andrew et al., 2012].

**Figure 1.**
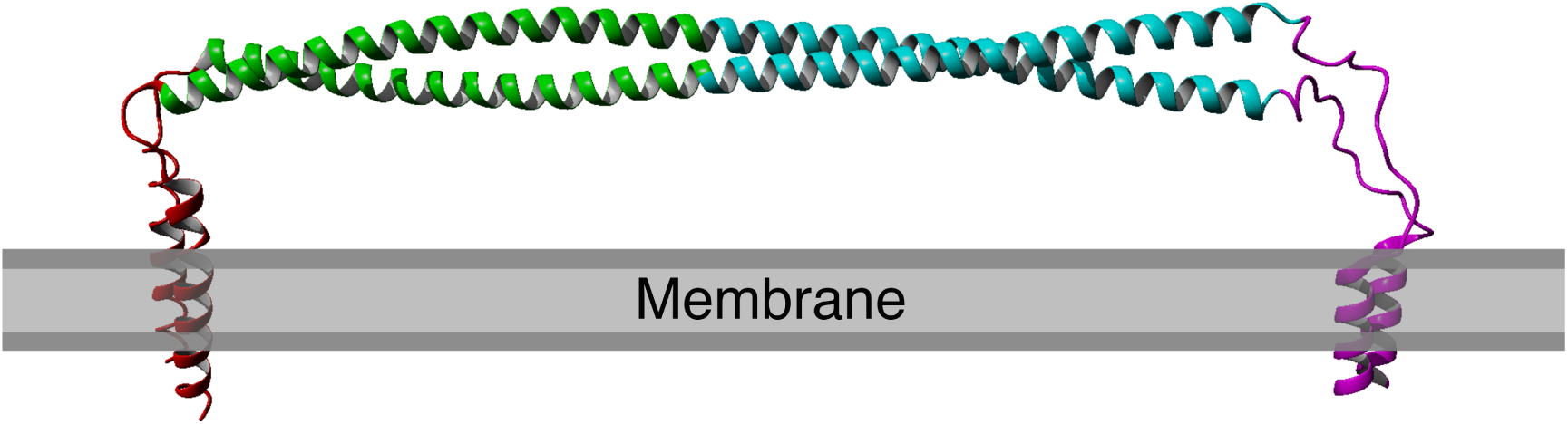
Structure of human Tetherin bound to the lipid membrane. Membrane anchors are shown in red (N) or magenta (C). The coiled-coil region within the ectodomain is shown in cyan. Image derived from the model produced by Ozcan and Berndsen, 2017

A few previous studies of Tetherin structure and function have used either scanning mutagenesis or analyzed a subset of known human mutations [Welbourn et al., 2015, Hammonds et al., 2012, Yang et al., 2010]. Cysteine scanning mutagenesis within the ectodomain showed that mutations in the coiled-coil region altered Tetherin dimerization and interfered with function, while cysteine mutations outside of this region were often functional [Welbourn et al., 2015]. The new cysteines changed the orientation of the helices and lead to changes in the folded state of Tetherin. Alanine scanning mutagenesis via changing of the sequence four alanines at a time showed two areas of the Tetherin ectodomain were sensitive to quadruple alanine mutation, however most mutations had no apparent effect [Hammonds et al., 2012]. The likely cause of functional defects was a change in protein localization [Hammonds et al., 2012]. Mutations that reversed the charge of three to six surface amino acids in the N-terminal portion of the ectodomain did reduce viral tethering, however whether this was due to alterations in structure or virus binding was unclear [Yang et al., 2010]. Deletions within the ectodomain also showed defects in viral tethering [Yang et al., 2010]. However, a later study by Andrew and coworkers showed that deletions in the ectodomain that alter the register of the alpha helix are detrimental while mutants with truncations that maintain the orientation of the helices are functional [Andrew et al., 2012]. These previous examples suggest that the Tetherin sequence is amenable to change as long as the alpha helical structure of the ectodomain is maintained.

In order to more fully understand the structure-function connection in Tetherin, we analyzed structural models of Tetherin containing one of 82 single nucleotide polymorphisms listed in the Ensembl database [Aken et al., 2017]. In all cases, the mutations did not affect the global structure of Tetherin and instead affected local structure primarily at the ends of the ectodomain. Mutations that occurred at the interface of the Tetherin monomers induced greater changes in protein dynamics than those that occur on the solvent-facing surface of the ectodomain. These data further support the idea that mutations in Tetherin are tolerated provided that the alpha helices in the ectodomain remain intact.

## METHODS

### Data, code, and analysis files

All macro files, R scripts, and data files are available on the Open Science Framework page for this project. These files include analysis of each mutation in terms of dynamics, secondary structure, and other simulation parameters.

### Production of Tetherin mutants

Tetherin mutants were generated from the membrane bound model of Tetherin described by [Ozcan and Berndsen, 2017] using the list of human mutations found in Ensembl as of August 2018 [Aken et al., 2017]. We wrote a YASARA macro to individually swap amino acids and minimally energy minimize the resulting structure. All models were equilibrated for 10 to 20 nanoseconds in explicit water solvent and a DOPC bilayer using an AMBER14 forcefield within YASARA Structure [Case et al., 2014, Krieger and Vriend, 2015, Krieger et al., 2009]. Simulation conditions were 298 K, pH 7.4, 0.9% mass fraction NaCl, at a density of 0.997 g/mL, with a 12 Angstrom non-bonded interaction cutoff, and a time step of 2.5 femtoseconds.

### Analysis of Models

Mutant simulations were analyzed using existing macros within YASARA to calculate simulation energies, root mean square fluctuation (RMSF), and the secondary structure over time. Data for each mutation were summarized using Rmarkdown. Root mean square deviation values were calculated using the Align function in YASARA structure. Global analysis of data was performed in R using the components within the *tidyverse* package and the *cluster* package [Ross et al., 2017, Wickham, 2009, Maechler et al., 2018].

## 1 RESULTS

### 1.1 Human missense mutations are absent from beginning of coiled-coil

The structure of Tetherin has been determined however the connection between the crucial functions of Tetherin and this structure is less clear [Swiecki et al., 2011, Hinz et al., 2010, Schubert et al., 2010, Yang et al., 2010]. The primary sequence of Tetherin is not well-conserved across species, although the general organization appears to be [Blanco-Melo et al., 2016]. Areas of structural or functional importance generally have a conserved protein sequence. Given the lack of apparent sequence conservation between species, it is difficult to predict the important structural features of Tetherin. We decided to compare sequences within the human species to identify areas of amino acid stability and therefore relative importance. We retrieved variation data from the Ensembl database containing the missense mutations within the ectodomain of Tetherin. Mapping the amino acid locations of SNPs showed that there was a lower density of mutations between residues 93 and 117 (Figure 2A). We note that synonymous mutations, which do not affect the amino acid sequence, still occur in this region, but amino acid changes appear to be lacking. Although the data set is limited in scope, the data suggest that the amino acids in this region, which includes one disulfide and the beginning of the coiled-coil region of the ectodomain, are important for Tetherin structure or function. Additionally, we mapped the mutations on the 3MQB structure and observed that 11 of 78 mutations appear to occur on the exterior of the protein (“outside”) rather than at the interface of the Tetherin dimer (“inside”) (Figure 2B) [Yang et al., 2010]. This distribution supports the previously demonstrated role of dimerization in viral tethering and stability [Andrew et al., 2009, Schubert et al., 2010, Welbourn et al., 2015, Du Pont et al., 2016]. We also observed that the areas of highest mutation density were in the N-terminal portion of the ectodomain, which has been suggested to be more flexible and is stabilized by the inter-molecular disulfides [Welbourn et al., 2015, Du Pont et al., 2016], or at the C-terminal end of the coiled-coil region, which in structural simulations unwinds relatively easily as it contains a non-canonical heptad [Andrew et al., 2012, Du Pont et al., 2016, Ozcan and Berndsen, 2017]. Therefore the locations of SNPs align well with previous functional and structural data on Tetherin. However, the potential effects of these mutations on Tetherin structure are unclear.

**Figure 2.**
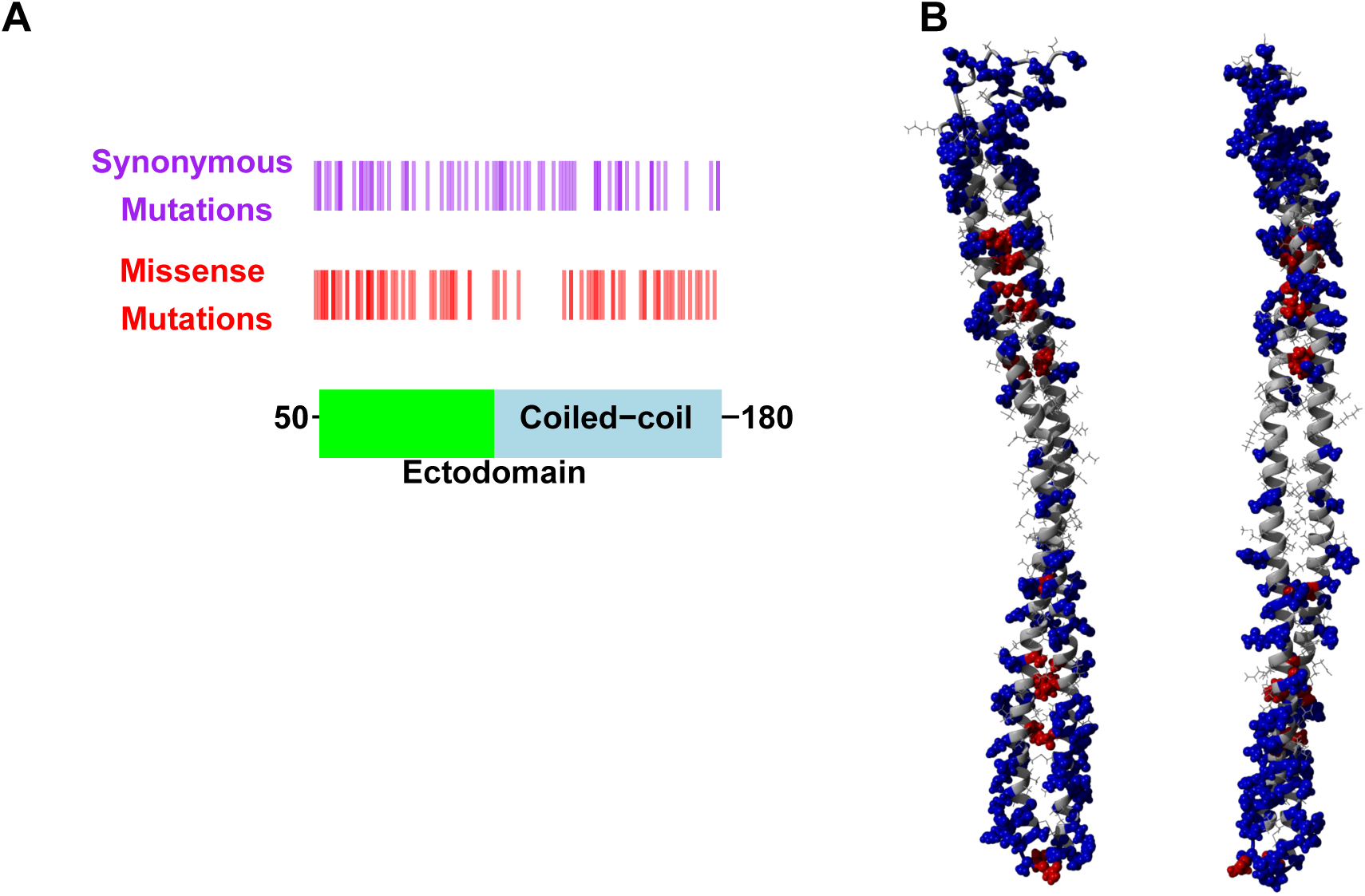
Human mutations within Tetherin (A) Cartoon showing domain organization and location of synonymous (purple) and missense mutations (red) in Tetherin (B) Model showing the location of the mutants on the inside (red) or outside (blue) of the Tetherin ectodomain.

### 1.2 Missense mutations do not alter global structure

To determine the association between structure and function in Tetherin mutants, we computationally made point mutations in the previously described full-length model of Tetherin [Ozcan and Berndsen, 2017]. All models equilibrated within 10 to 20 ns of simulation. We then compared the root mean square deviation, root mean square fluctuation, and secondary structure for all mutations to a similar simulation on a model lacking mutations to identify structural and dynamic changes due to the mutation. The root mean square deviation (RMSD) is a measure of the differences in the 3-D position of one or more elements of the ectodomain. We calculated the RMSD of the average structure from each mutant simulation from alignment of the ectodomain alone with a wild-type Tetherin that had also been equilibrated. The data were then plotted as RMSD vs. amino acid number as shown in Figure 3. Any trends of large increases in the RMSD value would suggest a strong effect on the global Tetherin structure. We observed that there were no trends in the RMSD values for the average structures across the ectodomain and no single mutation globally disrupted the coiled-coil Figure 3. These data suggest that mutations in Tetherin do not cause major changes to the helical structure.

**Figure 3.**
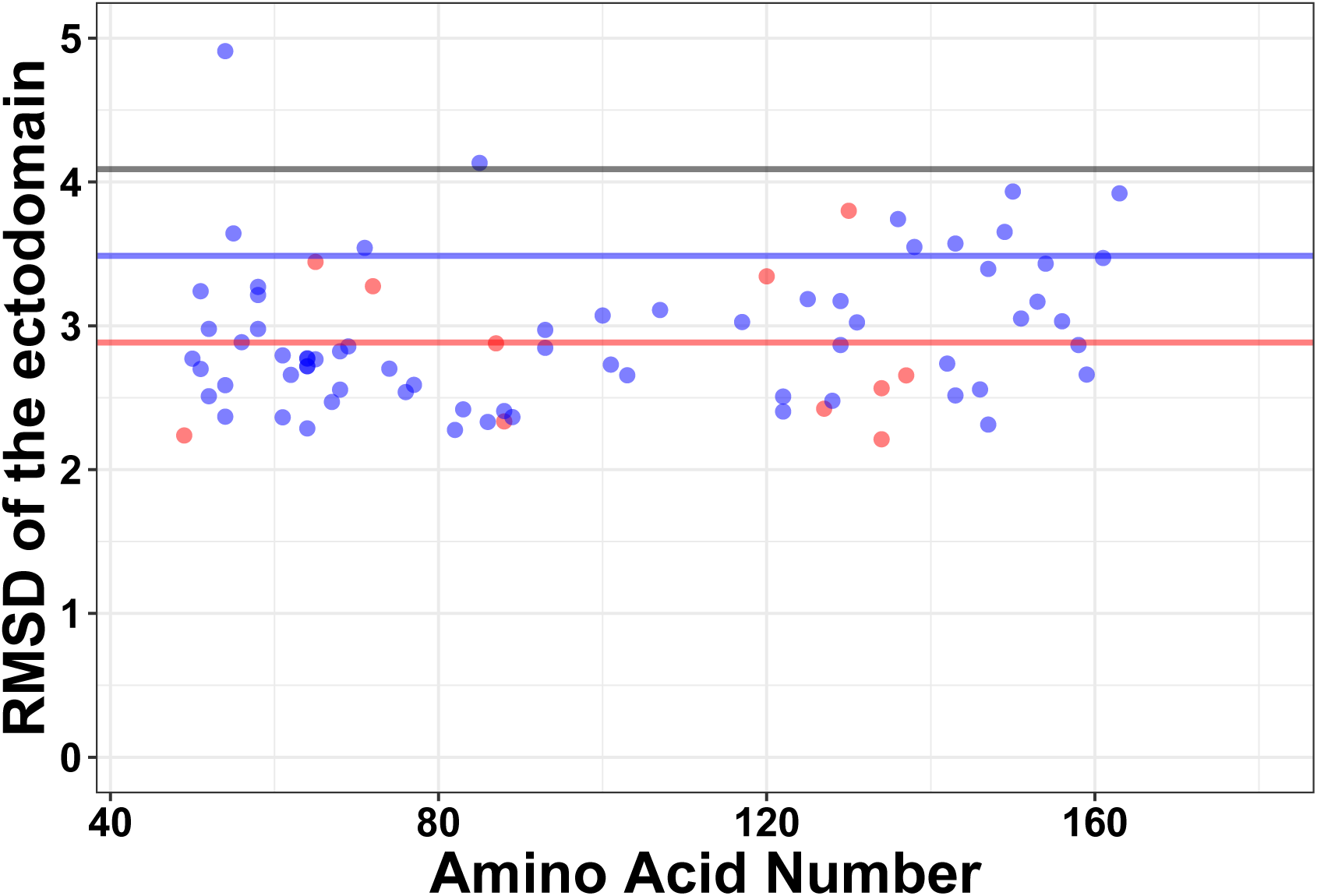
Few global changes in structure induced by mutations as shown by the root mean square deviation plotted against amino acid number. Red points are those positions that are found on the inside of the Tetherin dimer, while blue points are those positions on the outside of the dimer. Red, blue, and gray lines show the mean, mean + one and mean + two standard deviations of the RMSD values respectively

### 1.3 Mutations alter local Tetherin structure

Single changes to the amino acid sequence may not cause drastic changes to protein structure. Therefore, we next analyzed our simulations for local changes in helical structure. Based on the distance between hydrogen bond donors and acceptors, amino acid positions were binned into types of secondary structure at 400 steps throughout the simulation. Representative plots of secondary structure across the simulation are shown in Figure 4. Amino acids Ala100Pro and Ile120Phe showed distinct changes in secondary structure compared to the wild-type. Proline is well-known to be disruptive to alpha helices because it is unable to form the hydrogen bonds characteristic of this secondary structure [Pace and Scholtz, 1998]. Phenylalanine is aromatic and occupies a larger volume than isoleucine suggesting the local secondary structure changes are due to the alterations in the interface between the two helices. We note that none of the local changes in secondary structure spread along the coil. Instead, cracks in the helices stayed localized to a single area within the ectodomain, supporting the biochemical and computational findings of [Du Pont et al., 2016]. We next analyzed where the changes in structure were localized and plotted the number of times an amino acid adopted a different environment versus position (Figure 4). The effects on secondary structure appear primarily at the ends of the coil, rather than proximal to the site of substitution except for a few mutations (described below). Overall, these findings support the relative lack of effects on global helical structure that we observed as the ends of the coil are suggested to be unstructured leading to the terminal membrane anchors of the protein [Swiecki et al., 2011].

**Figure 4.**
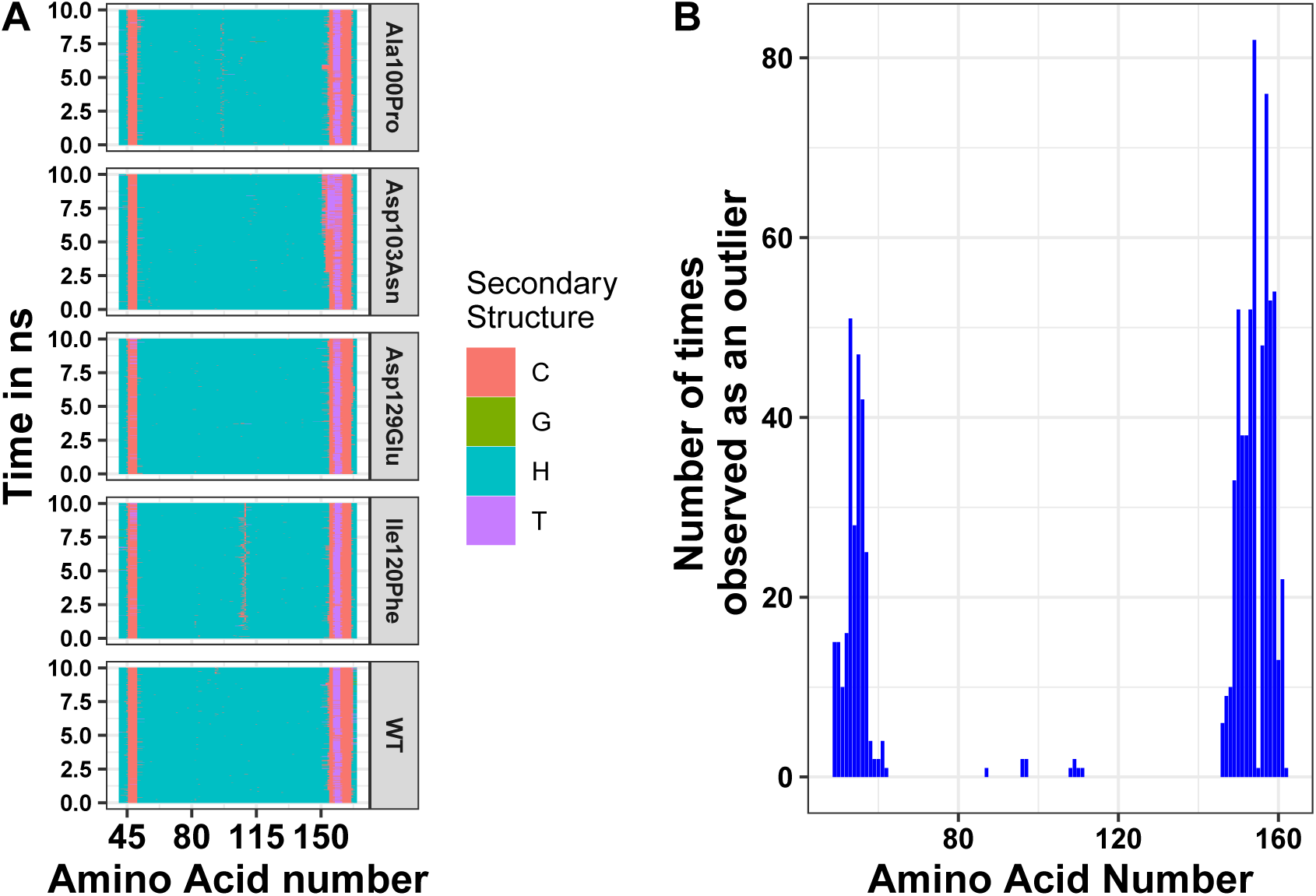
(A) Secondary Structure of BST-2 Mutant Simulations Over Time for Ala100Pro, Ile120Phe, Asp129Glu, and Asp103Asn. Secondary structures were assigned by YASARA based on hydrogen bonding distances and psi angles at 25 picosecond intervals. ‘H’ stands for alpha helix, ‘T’ for turn, ‘C’ for coil, ‘E’ for beta bridge, and ‘G’ for 3/10 helix. (B) Plot showing the number of times secondary structure was perturbed at each location for all simulations

While most mutations had minimal effect on the helical structure of Tetherin, five mutations did disrupt the helices. Gly64Pro and Ala100Pro induced alterations in secondary structure near to the site of the polymorphism. Asn65Asp and Thr67Ile cause changes at position 87 and 110/111 respectively, which are distant from the polymorphism site. In neither of these latter two groups do the alterations in structure include more than two consecutive amino acids. Ile120Phe however induces a disruption in secondary structure that is not directly adjacent to the mutation site and is maintained throughout the simulation. Visualization of the energy minimum structure from the simulation for both the wild-type and Ile120Phe structure, shows that Ile120 is an interior amino acid and the side chains could form a van der Waals contact(Figure 5A). However, in the structure of the Phe substitution, the side chain is displaced from the interface (Figure 5A). Further comparison of the distance between the Tetherin monomers and at sites along the helix shows that the Phe substitution noticeably changes distribution of measurements on either side of the mutation over a region 18 amino acids in length (Figure 5B). In comparison, the Ala100Pro mutation does not alter distribution of distances except for slight differences adjacent to the mutation site (Figure 5C). These data suggest that the structure of Tetherin tolerates local changes in orientation and configuration that do not alter the global structure of the protein.

**Figure 5.**
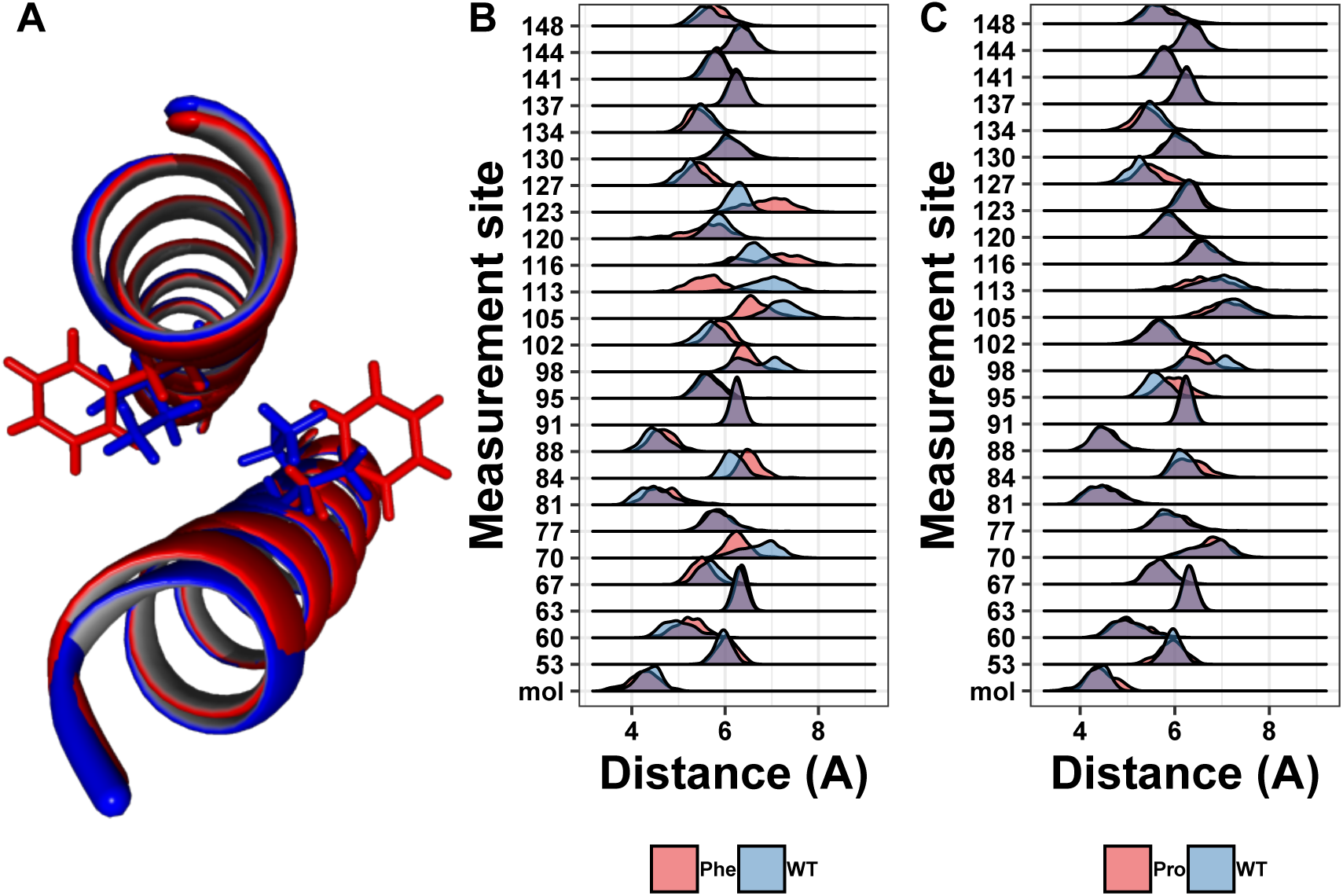
(A) Differences in side chain position between the wild-type Isoleucine (blue) and the mutation Phenylalanine (red) (B) Ridgeline plot showing the range of distances between the A and B molecules (mol) or the Calpha at the indicated positions comparing the Ile120Phe simulation (red) and the wild-type simulation (blue). Overlap is shown in purple. (C) Ridgeline plot showing the range of distances between the A and B molecules (mol) or the Calpha at the indicated positions comparing the Ala100Pro simulation (red) and the wild-type simulation (blue). Overlap between the two conditions is indicated by purple coloring.

### 1.4 Effect of mutations on ectodomain dynamics

Protein structure is not just based on the static arrangement of amino acids, but the ensemble of structures that the protein can adopt, which is sometimes referred to as protein dynamics. To quantify dynamics, we analyzed the root mean square fluctuation (RMSF), which describes the amount of movement each amino acid undergoes during the simulation. We then correlated the per residue RMSF values between each mutation simulation and the wild-type simulation. We observed moderate correlations at most positions but wanted additional methods to analyze the shape of the cluster. Using the central point in the cluster would allow for easier visualization of how each mutation affects the global dynamics of the system and allow for easier comparison of the effects of mutations. Therefore, we next calculated the mean and medoid centers of the RMSF clusters. The data for the medoid and mean cluster centers in the mutants correlated well suggesting that our data are normally distributed (Figure 6). We plotted all the data determined by the k-means method at each amino acid position to determine if there were any trends or clustering. As seen in the right pane of Figure 6, mutations at the ends of the ectodomain tended to have larger k-mean values suggesting larger RMSF values and more dynamics in the simulation. Mutations on the interface of the two helices did not increase dynamics relative to the outside mutations. These data are consistent with our previous finding that disruptions to secondary structure occurred preferentially toward the termini of the ectodomain and that the ectodomain can tolerate some changes to the structure.

**Figure 6.**
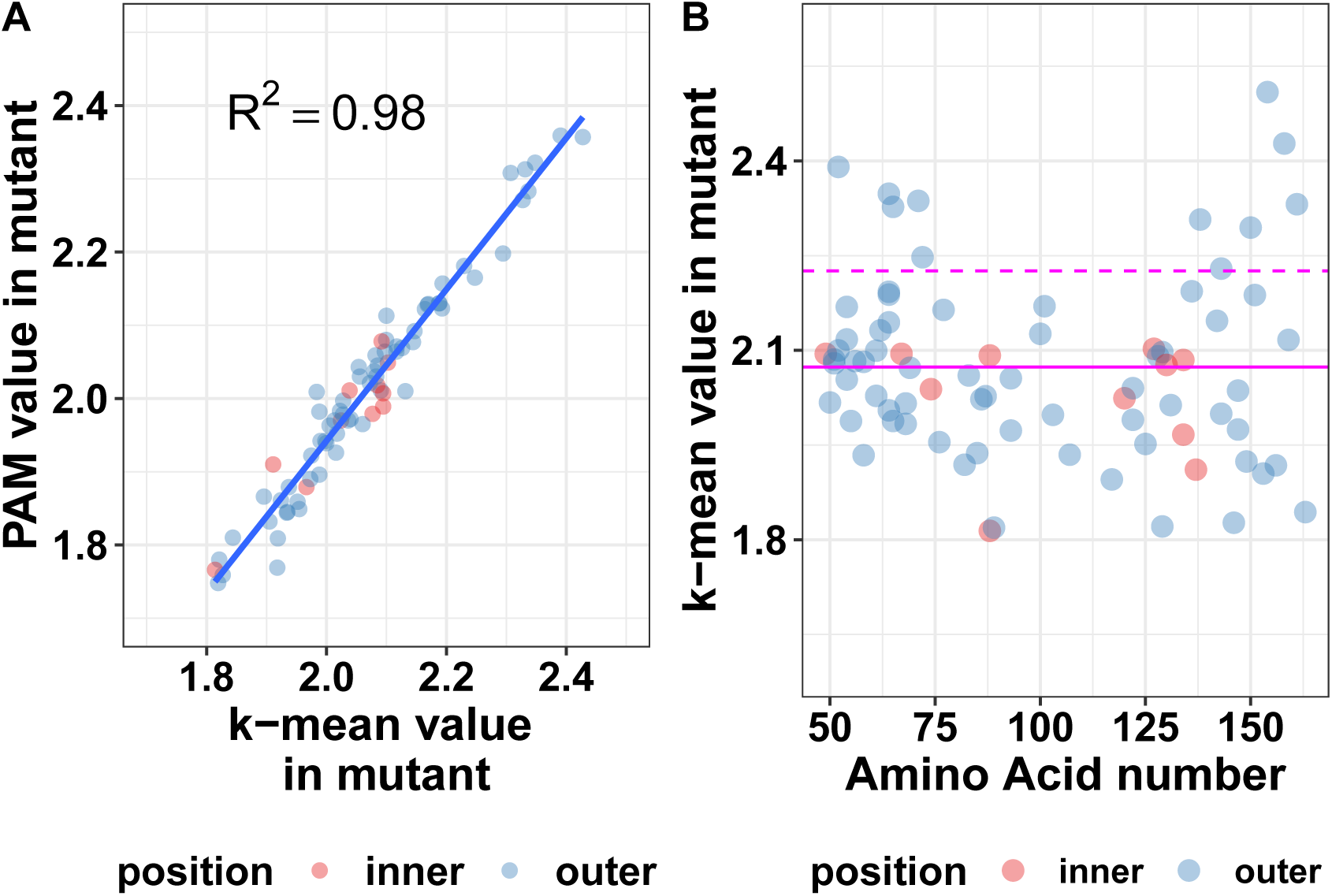
Mutation effects on protein dynamics (A) correlation of k-means and PAM values (B) scatterplot of k-mean values at each amino acid position. Magenta lines indicate the mean value (solid) and the mean plus one standard deviation (dashed) of the k-means has no known enzymatic functions and the interactions of Tetherin with viruses like HIV-1 which can rapidly adapt to immune system, this sequence flexibility is likely advantageous.

## DISCUSSION

The connection between Tetherin structure and function continues to be a significant question in the field. Putative orthologs of Tetherin show little sequence similarity and there are only structures for two orthologs [Hinz et al., 2010, Swiecki et al., 2011, Schubert et al., 2010, Yang et al., 2010, Blanco-Melo et al., 2016]. In order to gain more insight into the structural pliability of Tetherin, we analyzed the location and effects of Tetherin mutations found in the human genome. While the data set that we analyzed only includes data from those genome studies that deposit in Ensembl, this is the most comprehensive set of genomes available. Initial analysis showed that relative lack of missense mutations between amino acids 93 and 117. This section of Tetherin includes one cysteine which can form a disulfide, a glycosylation site, and also the first three heptads of the coiled-coil [Andrew et al., 2009, 2012]. While there are synonymous sequence changes in the DNA sequence along the entire length of the protein, this section lacks mutations that change the protein sequence. Our analysis of these mutations supports previous work showing this region as being important for flexibility or hinge motions [Hinz et al., 2010, Andrew et al., 2012, Ozcan and Berndsen, 2017]. Collectively, these data suggest that the flexibility of this region is tuned for a specific structure and/or function. Previous work suggested that this purpose could be to “break” in a controlled fashion during the transition from the membrane bound to the cell-virus bridging form [Ozcan and Berndsen, 2017]. However, more information is needed on the molecular functions of Tetherin to explore and test this idea fully.

Mapping of the mutations onto the structure of Tetherin [Yang et al., 2010] showed that most mutations occur on amino acids on the exterior of the helices rather than at positions at the dimer interface (Figure 2). The functional importance of Tetherin dimerization is well-established therefore, fewer interior mutations was not surprising. Throughout our analysis, mutations at the interface of the dimer showed similar behavior to the mutations on the outside of the dimer, suggesting disruptive mutations are not tolerated. One mutation, Ile120Phe, showed changes to the helical structure of the protein (Figure 4) which we linked to displacement of the side chain leading to changes in the distance between amino acids in each monomer of the dimer (Figure 5). The apparent disruption of the Tetherin dimer interface and relative change in the orientation of the monomers however was limited to a region local to the mutation site. These data suggest that the Tetherin dimer limits disruptions to localized regions and that more drastic reorganizations of the dimer are required, such as cysteine scanning mutagenesis performed by Welbourn et al. [2015].

In addition to the genetic stability of the middle section of Tetherin, we observed that most mutations had little if any effect on the overall structure of Tetherin. Calculation of the RMSD between the average structure during simulations and the wild-type protein showed no trends or large changes in the structure. Supporting these data were analysis of the secondary structure at each amino acid which showed most mutations appears to affect the ends of the ectodomain, which are known to be largely unstructured [Hinz et al., 2010, Andrew et al., 2012, Ozcan and Berndsen, 2017, Swiecki et al., 2011, Schubert et al., 2010, Yang et al., 2010]. Given the importance of the alpha helical structure of Tetherin to tensile strength, these findings are not surprising, although certainly unexpected given the number and diversity of mutations analyzed [Du Pont et al., 2016, Andrew et al., 2012]. This is most apparent in the Ala100Pro mutation, which we expected would cause drastic changes in Tetherin structure and dynamics. While the proline mutation does cause local changes in secondary structure (Figure 4), the effects of this mutation are localized to a few amino acids proximal to the mutation site. The detrimental effects of proline in helices are well-known Pace and Scholtz [1998], however the strength of the coiled-coil and the disulfides appears to prevent spreading of this “crack” in the helix [Du Pont et al., 2016]. Previously, Sauter and coworkers analyzed the functional effects of seven rare mutations in Tetherin from the human genome [Sauter et al., 2013]. For the mutations located in the ectodomain, most mutations had no effect on function [Sauter et al., 2013]. When we searched ClinVar immediately prior to submission of this paper for disease causing mutations in Tetherin, we did not find any single nucleotide polymorphisms in the ectodomain with known links to diseases or pathogenic phenotypes. These data lead to the idea that mutations are tolerated within Tetherin as long as they do not affect the overall helical structure and dimerization. Given that Tetherin

## ACKNOWLEDGMENTS

We thank the James Madison University Department of Chemistry and Biochemistry, Yasmeen Shorish, Kadir A. Ozcan, and the 4 – VA Organization for the resources, professional support, and models used throughout the project. We also thank the CHEM 361 students in Biochemistry 1 during Fall 2017 for their semester long work which helped shape the direction of the investigation. This work was supported in part by NSF-REU CHE-1461175. All data and files can be found on the Open Science Framework page for this paper at https://osf.io/hwkj4/.

